# Glucose availability tunes latent CD8+ T cell expansion potential through a mitogen-independent, mTOR-dependent regulatory switch

**DOI:** 10.64898/2026.01.16.699963

**Authors:** Tran Ngoc Van Le, Krittin Trihemasava, Lucien Turner, Kelly Rome, Aaron Wu, Will Bailis

## Abstract

Mitogenic signals are understood to license cell cycle progression and the metabolic reprogramming required for cell division, with acquired nutrients serving as permissive substrates. Here, we show that nutrient availability instead functions as a mitogen-independent regulatory input that dynamically controls CD8+ T cell proliferative potential. Activating stimuli have been shown to set T cell expansion capacity through their control of c-Myc expression, with the rate of c-Myc decay functioning as a division timer. We demonstrate that nutrient availability is sufficient to control c-Myc expression dynamics and dictates how division potential is stored and later actualized. Glucose-restricted T cells sustain proliferative potential and exhibit high AKT and ERK phosphorylation, despite limited growth. Upon glucose restoration, these cells rapidly increase c-Myc expression, accelerate through the cell cycle, and return to the expansion potential of glucose-replete controls, even after days of enforced restriction. Glucose restriction thus maintains a latent metabolic and mitogenic signaling state that is rapidly realized upon recovery. Mechanistically, mTOR signaling is required for this glucose recovery-driven proliferation, despite c-Myc and pERK remaining elevated following mTOR inhibition, indicating that glucose and mitogen signals operate through parallel rather than hierarchical control points. Altogether, these findings reveal that nutrient availability is not merely rate-limiting for proliferation but dictates the kinetics at which mitogenic signals are dissipated and realized. While mitogenic and nutrient cues converge on a shared anabolic network, they operate through distinct regulatory arms to coordinate the tempo and magnitude of clonal expansion, with implications for protective immunity and immunotherapy.

**Significance Statement:** CD8+ T cells rapidly proliferate to fight infections and cancer, often in variable nutrient environments. Activation signals are understood to control T cell expansion potential by setting c-Myc expression and its subsequent decay, with nutrients providing fuel. Here we find that glucose availability functions as an independent regulatory switch. Glucose-restricted T cells remain proliferatively poised for days, keeping pro-growth signaling and metabolic capacity primed. Upon glucose restoration, cells undergo a proliferative burst and catch up to glucose-replete counterparts. Although c-Myc expression rises upon glucose restoration, accelerated division kinetics instead require mTOR activity. These findings reveal that nutrient availability operates in parallel with mitogenic signaling, tuning the rate at which T cells store and realize their expansion potential.

## Introduction

Cellular proliferation is a fundamental process of primary mammalian cells that requires the integration of mitogenic signals and nutrient availability. Unlike unicellular organisms, which directly sense and scavenge nutrients from the environment and can adjust their proliferative rate strictly based on nutrient abundance, non-transformed mammalian cells rely on extracellular growth factors to instruct nutrient uptake and anabolic metabolism^1,2^. In the absence of a mitogenic signal, even in nutrient-replete environments, primary mammalian cells remain quiescent or undergo apoptosis, highlighting that nutrients alone are insufficient to drive proliferation^3–5^. In multicellular organisms, this dependence on mitogenic cues allows tissue-level coordination, ensuring that individual cells divide only when it is appropriate for the organismal homeostasis. Although growth factor stimulation under nutrient-limited conditions can transiently activate anabolic metabolism, sustained proliferation requires sufficient nutrient availability to maintain biosynthetic and bioenergetic demands^6–8^. Through this hierarchical relationship, growth factors act as instructive signals that license nutrient uptake and utilization, whereas nutrients provide essential substrates to support cell proliferation. Despite this established framework, it remains unclear whether proliferative potential, which refers to the number of divisions a clonal cell lineage can undergo, is predetermined by the initial mitogenic signal or dynamically regulated by subsequent modulation in nutrient availability, beyond that of a rate-limiting substrate.

Mitogenic cues engage several conserved and interconnected signaling pathways. These include the phosphoinositide 3-kinase (PI3K)-AKT-mechanistic target of rapamycin (mTOR) axis and the mitogen-activated protein kinase (MAPK)-extracellular signal-regulated kinase (ERK) cascade, which collectively regulate cellular growth, metabolism and proliferation. Within the PI3K axis, mTOR complex 1 (mTORC1) functions as a central metabolic sensor that integrates inputs from growth factors and nutrients to promote protein and nucleotide synthesis and cell cycle progression through the phosphorylation of p70 ribosome S6 Kinase (S6K) and eIF4E Binding Protein 1(4E-BP1)^9–16^. The role for mTOR complex 2 (mTORC2) is less well understood, but it is known to phosphorylate AKT at Ser473, reinforcing cell survival and glycolytic metabolism^17–21^. In parallel, activated ERK phosphorylates nearby cytosolic proteins and then translocates to the nucleus, where it regulates transcription factors controlling G1/S phase progression^22–27^. Although mTOR and ERK signaling coordinate multiple downstream targets, one key intersection is the transcription factor c-Myc. Once induced, c-Myc drives the expression of genes involved in glycolysis, amino acid transport and nucleotide biosynthesis, thereby coupling mitogenic signaling to the metabolic programs required for cell growth^28–31^. Despite their concerted actions in supporting proliferation, the requirements of mTOR and ERK in regulating c-Myc expression under various nutrient conditions are currently underexplored.

Protective adaptive immune responses require clonal expansion of antigen-specific CD8+ T cells^32,33^. CD8+ T cell activation and subsequent proliferation are initiated through three core inputs: T cell receptor (TCR) stimulation, co-stimulatory molecules and cytokines, which converge on shared downstream growth pathways such as PI3K-AKT-mTOR and MAPK-ERK to drive biomass accumulation, metabolic rewiring and cell cycle entry^34–41^. During the initial stages of activation, mTORC1 activity is essential for quiescence exit and the anabolic reprogramming necessary to support growth^42–48^. This early mTORC1-dependent phase also governs c-Myc expression as loss of Raptor, a core component of mTORC1, results in diminished c-Myc protein levels and disrupts the metabolic remodeling required for cell growth^31,43,49,50^. Importantly, c-Myc does not merely permit cell division, it sets CD8+ T cell proliferative potential. c-Myc expression scales with the strength and duration of TCR stimulation and gradually decays over time, functioning as a division timer as proliferation ceases following the loss of c-Myc^51^. Once cells have entered this c-Myc-dependent proliferative phase of activation, cell division is understood to become largely mTORC1-independent^42,43,52^.

While required, mitogenic signaling alone is not sufficient to sustain proliferation. Nutrient availability is well recognized as both rate-limiting and a key regulator of mTORC1 and c-Myc activity. Activation under glucose or amino acid restricted conditions both impairs proliferation as well as mTORC1 signaling and c-Myc expression^31,53–56^. This nutrient sensitivity is understood to contribute to CD8+ T cell dysfunction in nutrient-poor environments such as tumors, infected tissues, or inflamed sites, where glucose or amino acids can become restricted^31,54–58^. Although early activating signals establish the transcriptional and metabolic programs required for proliferation, it is unclear whether this proliferative potential is fixed at the moment of activation and only decays over time, or if the nutrient environment can influence the rate at which this potential is realized and/or stored^38,51,59^. Understanding how CD8+ T cell proliferative potential is differentially influenced by mitogen and nutrient availability is central not only to basic cell biology but also to therapeutic strategies aimed at enhancing CD8+ T cell expansion in metabolically deprived environments.

In this study, we set out to determine whether nutrient availability primarily acts as a rate-limiting factor or can independently modulate proliferative potential. We demonstrate that glucose-restricted CD8+ T cells maintain a latent metabolic and mitogenic signaling state, which enables the full recovery of expansion potential upon glucose restoration days later, without requiring additional TCR stimulation. Glucose-restricted CD8+ T cells maintain high levels of phospho-AKT(Ser473) and phospho-ERK, as well as elevated metabolic potential. Even after several days of restriction, restoring environmental glucose is sufficient to achieve the same level of expansion as cells maintained in glucose-replete conditions, without additional TCR or costimulation. While we observe a time-dependent decay of c-Myc protein in both restricted and replete conditions, glucose recovery triggers a rapid elevation in c-Myc expression and mitogenic signaling activity coinciding with proliferative recovery. Mechanistically, we demonstrate that mTOR signaling is required for glucose restoration-induced proliferation and uniquely regulates a suite of key transcripts governing cellular metabolism, despite c-Myc and phospho-ERK levels remaining high in mTORC-inhibited cells. These data indicate mTOR signaling is specifically required at points of cell cycle acceleration in CD8+ T cells, rather than cell cycle entry alone, operating in parallel to the monotonic role of c-Myc in division kinetics. Altogether, we reveal that rather than mitogenic signaling acting only upstream of environmental nutrient uptake to control T cell expansion, nutrient availability additionally functions as an independent signal that dynamically modulates how CD8+ T cell proliferative potential is stored and dissipated. These findings support a model in which nutrient availability can throttle expansion kinetics as CD8+ T cells navigate changing metabolic landscapes while retaining information from the initial antigen encounter, allowing them to flexibly execute their effector function across diverse disease contexts.

## Results

### CD8+ T cell expansion capacity is controlled by both the magnitude of TCR stimulation and nutrient availability

To investigate how mitogenic signaling and nutrient availability orchestrate CD8+ T cell expansion potential, we first sought to establish how these parameters control activation kinetics and proliferation in the same system. To examine how TCR strength influences CD8+ T cell proliferation, we used the OT-I TCR transgenic model, in which all T cells express a TCR that recognizes the high-affinity peptide SIINFEKL (N4) and the lower-affinity variant SIIGFEKL (G4)^60^. As expected, OT-I CD8+ T cells stimulated with either N4 or G4 or anti-CD3/anti-CD28 at various concentrations showed an affinity- and dose-dependent increase in total cell numbers (**Fig. 1A and S1A**)^37,38^. Stronger and higher-concentration TCR signals increased both the fraction of cells that became activated and the proportion that entered the cell cycle (**Fig. 1B, C and S1B-D)**. Consistently, the percentage of cells expressing c-Myc was elevated with increasing activating signal strength (**Fig. 1D and S1E)**^59^. Although the strength of the TCR stimulation also affected total c-Myc protein expression, these differences were less pronounced when quantified only within the c-Myc+ population (**Fig. S1F and G)**. Thus, TCR signal strength primarily regulates the frequency of c-Myc+ induction rather than the amount of c-Myc per individual c-Myc+ cell, consistent with prior reports^59^. By 48 hours post-activation, we further observed that the frequency of cells that had undergone at least one division also corresponded with TCR stimulation strength (**Fig. 1E and S1H**). Analysis of the mean division number (MDN) revealed that, aside from the lowest G4 concentration, all peptide-stimulated groups exhibited similar MDN values from day 3 onward (**Fig. 1F and S1I)**. Likewise, anti-CD3/CD28 stimulation showed a comparable pattern, though lower antibody concentration yielded a modest reduction in MDN (**Fig. S1J)**. The remaining difference in cell number for low peptide affinity and antibody concentration could be attributed to the reduced CD8+ T cell viability rather than slower cell division in fully activated cells (**Fig. 1G and S1K)**. Together, these findings indicate that, in the presence of co-stimulation and IL-2, the magnitude of TCR signaling predominantly governs expansion potential through its impact on activation and cell cycle entry kinetics, balanced against the pro-survival signals it confers, in keeping with previous reports^38^.

**Figure 1:**
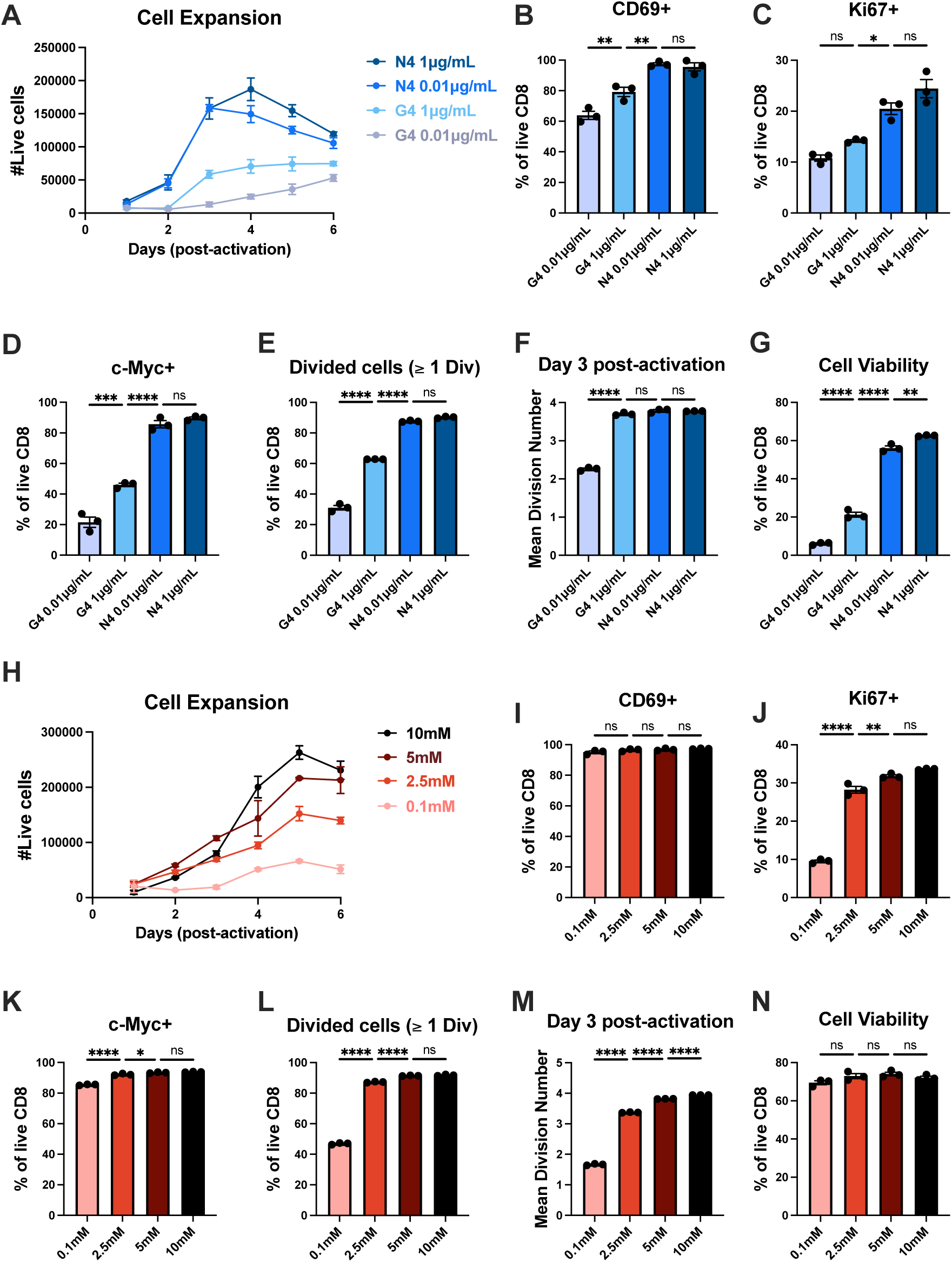
TCR stimulation strength and nutrient availability dictate T cell proliferation capacity through distinct mechanisms. CD8+ T cells were purified from OT-I spleen and lymph nodes that had been stimulated for 24 hours with either 1µg/mL or 0.01µg/mL of SIINFEKL (N4) or SIIGFEKL (G4) peptides and 2.5µg/mL anti-CD28. (**A**) Quantification of the number of live CD8+ T cells activated by different TCR affinity and concentration. (**B**) Frequency of CD8+ T cells expressing activation marker CD69 24 hours post-activation. (**C**) Frequency of CD8+ T cells entering cell cycle (Ki67+) 24 hours post-activation. (**D**) Frequency of CD8+ T cells expressing c-Myc 24 hours post-activation. (**E**) Frequency of divided cells (≥ 1 division) on day 2 post-activation. (**F**) Mean division number (MDN) on day 3 post-activation. (**G**) Frequency of live CD8+ T cells 24 hours post-activation. CD8+ T cells were purified from OT-I spleen and lymph nodes that had been stimulated for 24 hours with 1µg/mL of N4 peptide and 2.5µg/mL anti-CD28 in various glucose concentrations (10mM, 5mM, 2.5mM and 0.1mM). (**H**) Quantification of the number of live CD8+ T cells activated in different glucose concentrations. (**I**) Frequency of CD8+ T cells expressing activation marker CD69 24 hours post-activation. (**J**) Frequency of CD8+ T cells entering cell cycle (Ki67+) 24 hours post-activation. (**K**) Frequency of CD8+ T cells expressing c-Myc 24 hours post-activation. (**L**) Frequency of divided cells (≥ 1 division) on day 2 post-activation. (**M**) MDN on day 3 post-activation. (**N**) Frequency of live CD8+ T cells 24 hours post-activation. All error bars are representative of 3 technical replicates. Statistical significance was calculated using one-way ANOVA with multiple comparisons and Tukey’s correction.

To investigate the role of nutrient availability in CD8+ T cell proliferation, OT-I CD8+ T cells were stimulated with N4 peptide in media containing varying glucose concentrations, titrated from the standard RPMI-1640 concentration (10mM). As shown previously, we observed that the magnitude of CD8+ T cell expansion was proportional to the environmental glucose supplied (**Fig. 1H)**^55,56^. TCR-mediated activation appeared largely unaffected, as all N4-stimulated cells expressed comparable levels of the activation marker CD69 regardless of glucose levels (**Fig. 1I**). In contrast, glucose availability markedly influenced the rate of cell cycle entry. Despite exhibiting comparable c-Myc protein expression, cells cultured in higher glucose concentrations entered the cell cycle more efficiently (**Fig. 1J, K and S1L**). Correspondingly, the initial fraction of divided cells and MDN values scaled proportionally with glucose availability, suggesting that c-Myc-independent factors also modulate the division timer in response to nutrient availability (**Fig. 1L, M and S1M)**. In contrast to TCR signaling, environmental glucose had a minimal impact on CD8+ T cell viability (**Fig. 1N)**. Altogether, these data indicate that while both TCR signaling and glucose availability regulate CD8+ T cell expansion potential, each contributes through distinct mechanisms. Whereas the strength of TCR signaling controls the kinetics of activation, c-Myc expression, cell cycle entry, and death, glucose availability impacts cell cycle entry and division dynamics independent of activation, death, or c-Myc expression. Thus, mitogenic signaling and environmental glucose act as discrete inputs that operate through distinct mechanisms yet converge upon CD8+ T cell expansion capacity.

### CD8+ T cells maintained proliferative potential during glucose restriction

Prior studies have shown that activating signal strength sets the long-term T cell clonal expansion capacity^38^. Finding that TCR stimulation and glucose availability influence CD8+ T cell expansion potential in distinct ways, we next asked whether these two inputs could be uncoupled. To address this, we activated CD8+ T cells in either control (10 mM glucose) or glucose-restricted media (0.1 mM glucose) for 24 hours, after which activating stimuli were removed. We then allowed glucose-restricted cells to grow for 2, 3 or 4 days post-activation before restoring them to 10 mM glucose (**Fig. 2A)**. Regardless of when glucose was restored, previously glucose-restricted cells regained proliferative capacity and achieved comparable expansion to control cells (**Fig. 2B)**. By day 4, rescued cells exhibited a significantly greater number of total divisions compared to glucose-restricted cells (**Fig. 2C and D)**. Over the preceding day, glucose-restored cells underwent more divisions than controls **(Fig. 2E).** This reveals that proliferative potential is preserved during glucose restriction and can be reengaged once glucose becomes available.

**Figure 2:**
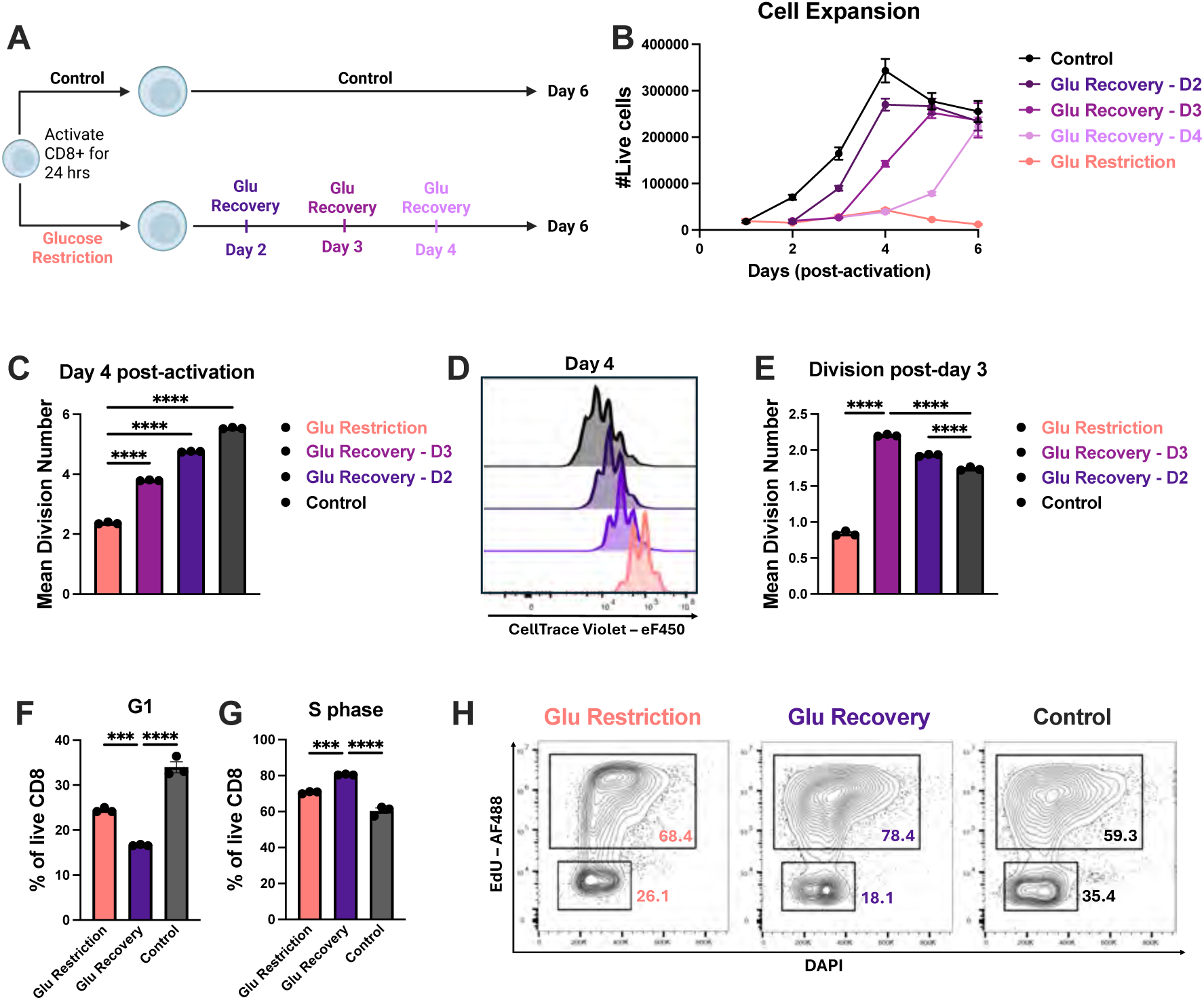
Maintenance of proliferative potential in CD8+ T cells during glucose restriction. (A) Experimental schematic of glucose restriction and recovery. Mouse CD8+ T cells were activated with 1µg/mL of N4 peptide and 2.5µg/mL anti-CD28 in either 10mM glucose (control) or 0.1mM glucose (glucose restriction). Cells were removed from stimulation after 24 hours. Glucose-restricted cells were restored with 10 mM glucose on either day 2, day 3, or day 4 post-activation. (**B**) Quantification of the number of live CD8+ T cells over time. (**C**) Mean division number (MDN) on day 4 post-activation. (**D**) Representative histogram of CellTrace Violet on day 4 post-activation. (**E**) Quantification of proliferative progression, represented as the increase of MDN from day 3 to day 4 post-activation, relative to the same cell population. Glucose-restricted, 24-hour glucose-recovered, and glucose-sufficient cells (on day 3 post-activation) were incubated with 5µM EdU for 4 hours, followed by Click-iT reaction for EdU detection and DAPI staining for cell cycle analysis. (**F**) Frequency of cells in G1 phase. (**G**) Frequency of cells in S phase. (**H**) Representative plot of EdU versus DAPI. All error bars are representative of 3 technical replicates. Statistical significance was calculated using one-way ANOVA with multiple comparisons and Tukey’s correction.

We next examined how glucose recovery influenced cell cycle kinetics. Glucose restoration led to a significant shift in cells transitioning from G1 to S phase, compared to both glucose-restricted and control conditions (**Fig. 2F-H)**. These data indicate that glucose restoration can tune the rate at which cells progress through the G1-S transition, resulting in the accelerated pace of cell division observed. Collectively, these data suggest that glucose restriction preserves a latent division potential rather than acting solely as a metabolic constraint. The rapid acceleration in cell cycle dynamics upon glucose restoration further indicates that a glucose-sensing mechanism likely mediates this switch-like behavior.

### Glucose restriction preserves CD8+ T cell metabolic potential, which is rapidly realized upon recovery

In light of this rapid change in division kinetics, we aimed to determine how glucose recovery reshapes CD8+ T cell metabolism. To address this, we evaluated the relative contribution of oxidative phosphorylation and glycolysis to total ATP production^61,62^. Glucose-restricted cells displayed significantly higher basal and maximal respiration compared to controls, resulting in an increased spare respiratory capacity (**Fig. 3A-E)**. In glucose-recovered cells, both respiratory capacity and glycolytic activity increased further, suggestive of metabolic reprogramming (**Fig. 3A-I)**^63^. Consistent with this elevated metabolic activity, ATP production derived from either glycolysis or mitochondrial respiration was markedly elevated in glucose-recovered cells (**Fig. 3J)**. Given these changes, we next examined whether glucose restoration impacted mitochondrial biology. Replenishing glucose to control levels resulted in a pronounced increase in both mitochondrial mass as well as membrane potential, reflecting the heightened respiratory capacity of these cells (**Fig. 3K and L)**. Our findings suggest that glucose restriction preserves CD8+ T cells in a metabolically poised state, which can be quickly tapped to fuel the accelerated division kinetics upon glucose recovery.

**Figure 3:**
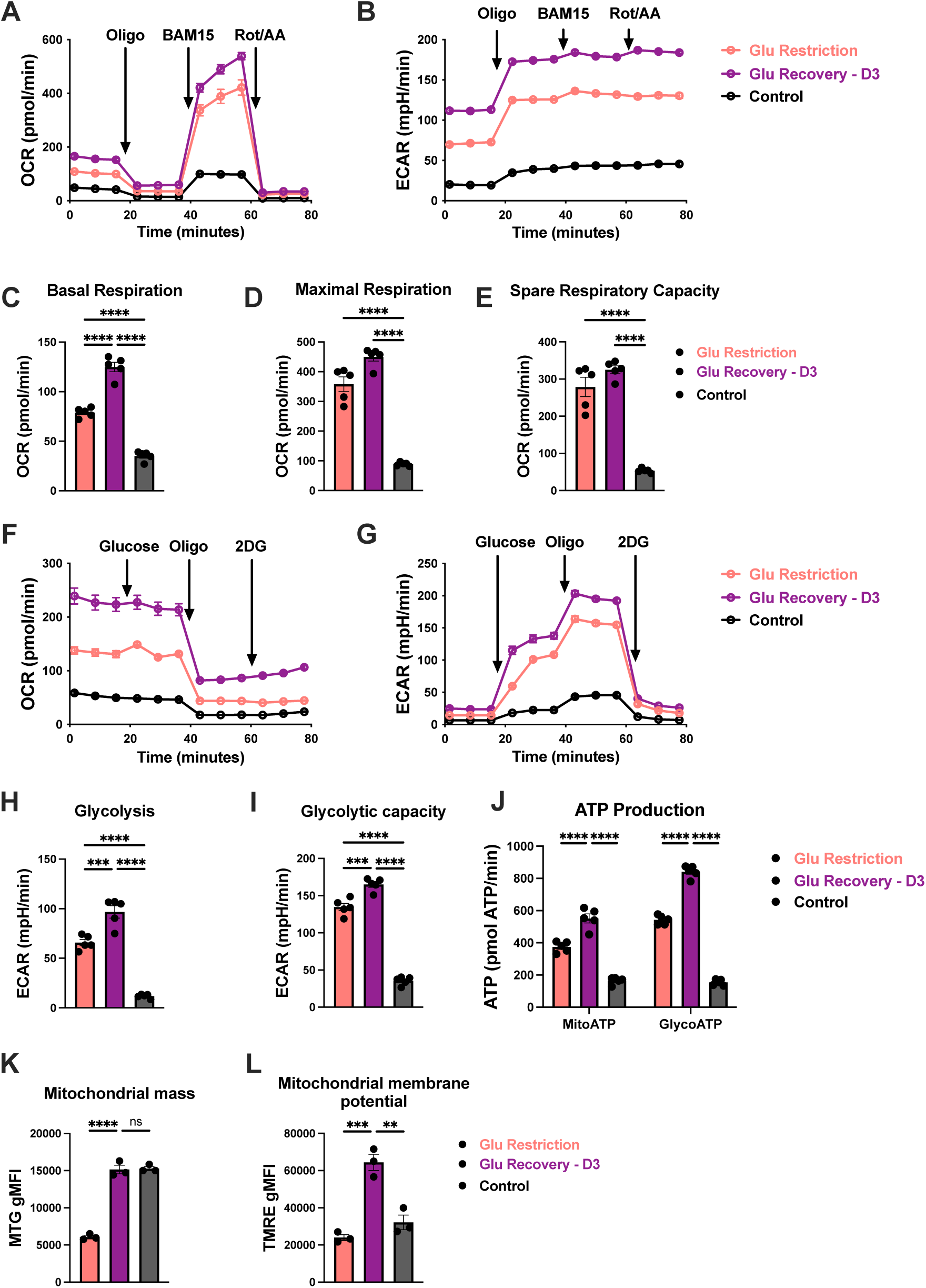
Glucose limitation maintains CD8+ T cell metabolic readiness for rapid enhancement upon glucose recovery. (**A**) Oxygen consumption rate (OCR) and **(B)** Extracellular acidification rate (ECAR) of glucose-restricted, glucose-recovered and control CD8+ T cells at baseline or in the presence of these subsequent drugs: oligomycin (ATP synthase inhibitor), BAM15 (uncoupling agent), and Rotenone/Antimycin A (complex I/III inhibitor). (**C**) Basal, **(D)** maximal respiration, and **(E)** spare respiratory capacity calculated from the mitochondrial stress test. (**F**) OCR and **(G)** ECAR of glucose-restricted, glucose-recovered and control CD8+ T cells at baseline or in the presence of these subsequent compounds: glucose, oligomycin and 2-Deoxy-D-glucose (2DG, glycolytic inhibitor). (**H**) Glycolytic basal rate and **(I)** glycolytic capacity measured from the glycolytic stress test. (**J**) ATP production was calculated from glycolysis and mitochondrial respiration. (**K**) Mitochondrial mass was measured from MitoTracker Green (MTG) geometric mean fluorescence intensity (gMFI). (**L**) Mitochondrial potential was measured from TMRE gMFI. All error bars are representative of 3-5 technical replicates. Statistical significance in (C, D, E, H, I, K and L) was calculated using one-way ANOVA with multiple comparisons and Tukey’s correction, and statistical significance in (J) was calculated using two-way ANOVA with multiple comparisons and Tukey’s correction.

### Glucose recovery re-engages mitogenic signaling without additional activating stimuli

TCR, costimulatory signal strength and cytokines together are understood to set the peak level of c-Myc induced in a population upon activation, after which c-Myc levels progressively decline, functioning as a temporal regulator of division potential^38,51,59,64^. Given that nutrient availability can recover division potential even after this temporal window has progressed, we asked whether glucose restoration bypasses the c-Myc division timer or actively “rewinds” it. To address this, we first evaluated the dynamics of mTORC, ERK, and c-Myc signaling over the course of activation and expansion in control and glucose-restricted cells. Glucose restriction resulted in reduced mTORC1 signaling, characterized by decreased total and phosphorylated S6K and S6, with 4E-BP1 expression remaining slightly elevated (**Fig. 4A and B)**. Consistent with the slower proliferation rates, we found that c-Myc expression was reduced in glucose-restricted cells, while decaying over time in both conditions (**Fig. 4C)**. In contrast, glucose-restricted cells displayed elevated phospho-AKT (Ser473) and phospho-ERK (Thr202/Tyr204), suggesting that limited glucose availability supported pathway activity (**Fig. 4D)**.

**Figure 4:**
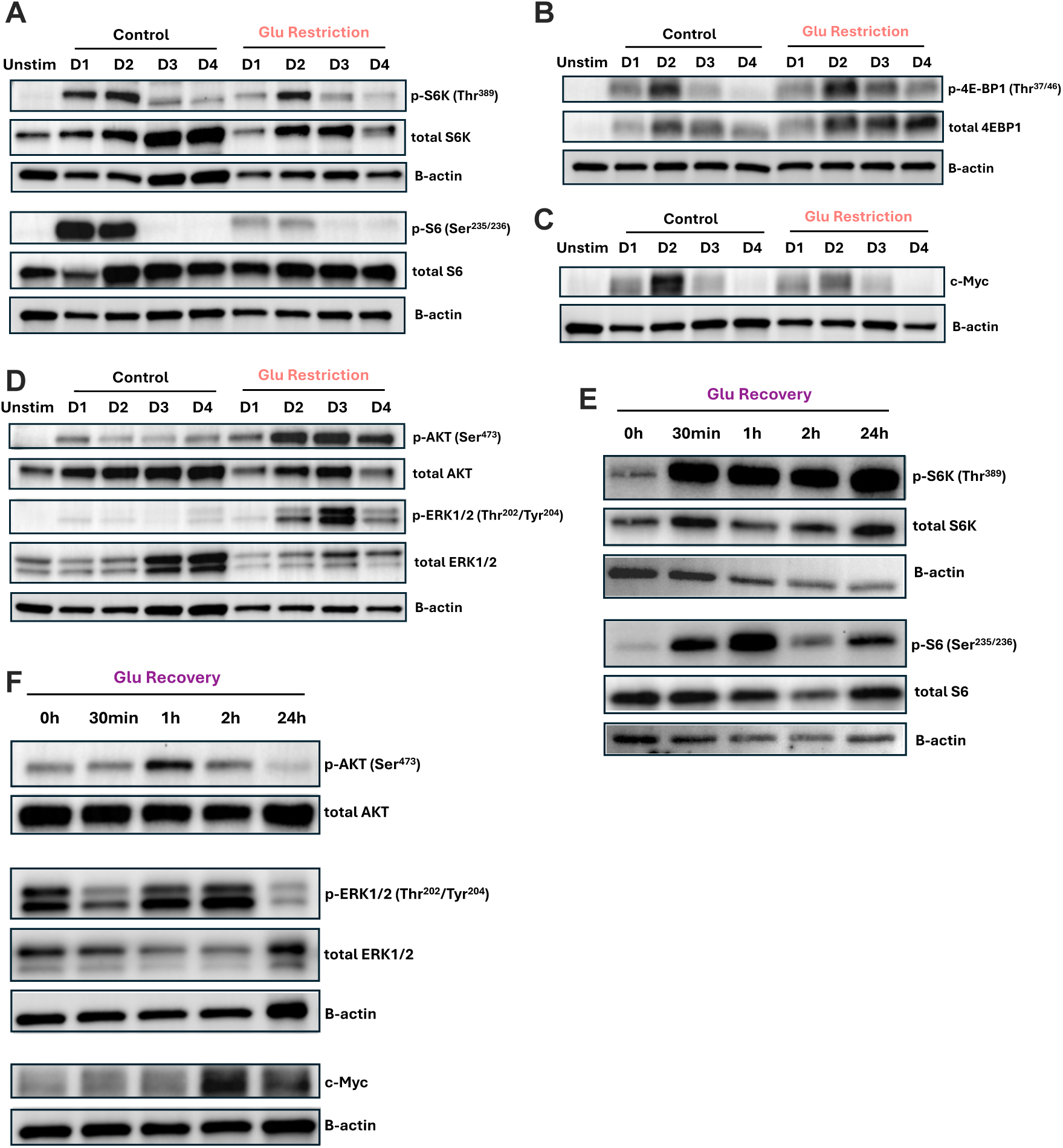
Glucose supplementation rapidly induces mitogenic pathways in the absence of additional activating inputs. Time course immunoblots across day 1-4 of control (10mM glucose) and glucose-restricted (0.1mM glucose) samples of (**A**) total and p-S6K (Thr389), p-S6 (Ser235/236), (B) total and p-4EBP1 (Thr37/46), (**C**) c-Myc, (**D**) total and p-AKT (Ser473), p-ERK1/2 (Thr202/Tyr204), with B-actin as loading control. Immunoblots of glucose-restricted cells at indicated time points after recovery on day 3 post-activation of (**E**) total and p-S6K (Thr389), p-S6 (Ser235/236), (**F**) total and p-AKT (Ser473), p-ERK1/2 (Thr202/Tyr204) and c-Myc, with B-actin as loading control.

We next sought to determine how glucose recovery impacted these signaling dynamics. Consistent with mTORC1’s known capacity to sense glucose availability, restoration led to rapid increases in S6K and S6 phosphorylation (**Fig. 4E)**^65–71^. This was accompanied by a further elevation of AKT (Ser473) and ERK (Thr202/Tyr204), indicating a broad reengagement of pro-growth signaling pathways (**Fig. 4F**). In parallel, c-Myc expression increased shortly following glucose recovery and remained elevated 24 hours later, coinciding with the accelerated proliferative response (**Fig. 4F**). Together, these data support a model in which glucose restriction preserves selective pro-growth signaling, and glucose recovery can re-license mitogenic pathways. Moreover, our data demonstrate that rather than being a fixed activation-regulated c-Myc division timer, environmental nutrients can independently enhance c-Myc expression and restore cell division potential.

### mTORC signaling is required for accelerated division and metabolic reprogramming upon glucose recovery

Thus far, we observed that glucose availability dynamically regulates multiple pro-growth signaling pathways, sustaining some during restriction (ERK, mTORC2-AKT) while rapidly re-engaging others after recovery (c-Myc, mTORC1). While c-Myc is the canonical regulator of sustained CD8+ T-cell proliferation following quiescence exit, we found that TCR and glucose availability differentially impact initial c-Myc expression, despite both affecting expansion potential (**Fig. 1D and K**). In contrast, mTOR is thought to play a lesser role in T cell proliferation beyond mTORC1’s requirement during quiescence exit, yet both mTORC1 and mTORC2 are highly responsive to glucose availability^42–44,52,65–70,72–74^. We therefore tested the hypothesis that mTOR signaling plays a unique role in promoting the accelerated division kinetics post-glucose restoration. To address this, we treated activated CD8+ T cells at the time of glucose recovery with either the dual mTORC1 and mTORC2 inhibitor Torin1, or rapamycin, which primarily targets mTORC1 but can reduce mTORC2 activity with prolonged exposure^75–77^. In keeping with this, both drugs suppressed the activity of mTORC1 and mTORC2 pathway targets across conditions (**Fig. 5A**). Although mTOR activity was reduced, the already limited proliferative capacity of glucose-restricted cells did not further decline in response to either inhibitor (**Fig. 5B**). In contrast, while control cells exhibited significant sensitivity to the mTOR pathway, the magnitude of this effect was minor compared to what we observed following glucose recovery. Indeed, mTOR inhibition abrogated the accelerated division kinetics found in glucose-recovered CD8+ T cells, highlighting the unique and heightened reliance of these cells on the mTOR pathway (**Fig. 5B**). Having seen this, we next asked whether mTOR signaling was necessary for the elevated phospho-ERK and c-Myc protein levels observed post-glucose recovery. Despite the pronounced effect of mTOR inhibition on proliferation, both c-Myc expression and ERK phosphorylation remained unchanged or were slightly elevated across conditions (**Fig. 5C**). These findings suggest that mTOR signaling is a principal driver of accelerated proliferation in glucose-recovered cells, acting independently of enhanced MAPK-ERK and c-Myc activity.

**Figure 5:**
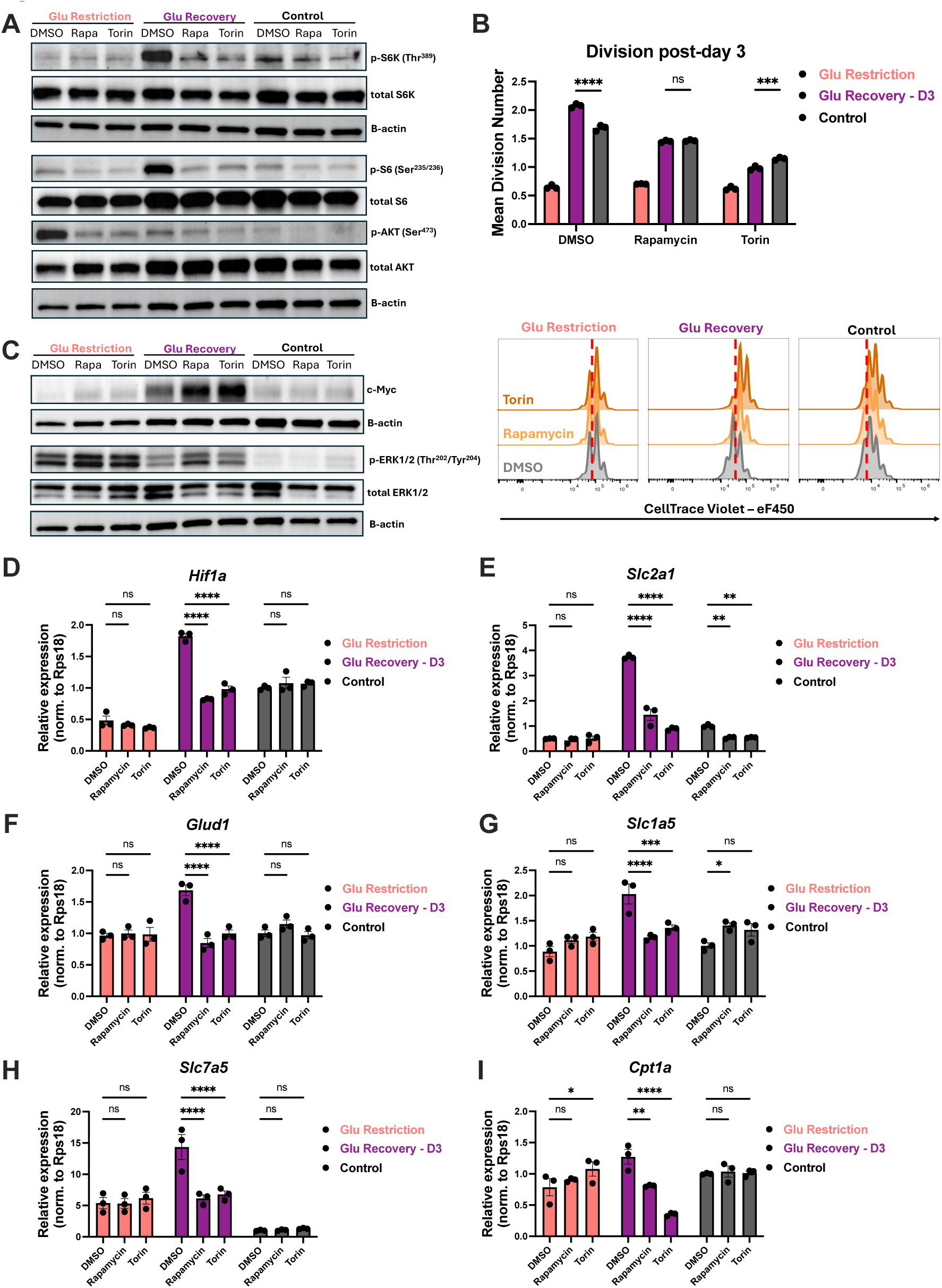
mTORC signaling drives accelerated proliferation and metabolic adaptations in glucose-recovered cells. On day 3 post-activation, cells were treated with DMSO, rapamycin (50nM), or Torin1 (50nM) within their corresponding media conditions for 24 hours. (**A**) Immunoblots of total and p-S6K (Thr389), p-S6 (Ser235/236), and p-AKT (Ser473), with B-actin as loading control. (**B**) Mean division number with representative histogram of CellTrace Violet. (**C**) Immunoblots of c-Myc and total and p-ERK1/2 (Thr202/Tyr204), with B-actin as loading control. Relative mRNA expression of (**D**) hypoxia inducible factor 1 subunit alpha (*Hif1a*), (**E**) glucose transporter Glut1 (*Slc2a1)*, (**F**) glutamate dehydrogenase 1 (*Glud1*), (**G**) neutral amino acid transporter (*Slc1a5*), (**H**) large neutral amino acid (*Slc7a5*) and (**I**) rate-limiting enzyme of fatty acid oxidation (*Cpt1a*). All error bars are representative of 3 technical replicates. Statistical significance was calculated using two-way ANOVA with multiple comparisons and Tukey’s correction.

We next aimed to test whether mTOR signaling has a unique gene regulatory role during glucose recovery. We found that glucose recovery alone led to a broad increase in the expression of genes involved in glycolysis (*Hk2, AldolA, Eno1, Pgk1, Ldha*), *Hif1a*, and *Slc2a1* (glucose transporter 1), consistent with the elevated glycolytic capacity of these cells (**Fig. 5D, E and S2A-E)**^53,78–80^. Additionally, glucose supplementation led to elevated glutamate dehydrogenase 1 (*Glud1*) transcript abundance as well as the glutamine and large neutral amino acid transporters, *Slc1a5* and *Slc7a5,* respectively (**Fig. 5F-H)**. While several mTOR pathway targets, particularly those involved in glycolysis, were sensitive to rapamycin and Torin1 in control cells, we identified a subset of genes that were selectively regulated upon glucose restoration. In particular, glucose-recovered cells showed a unique mTOR dependence for the expression of *Hif1a*, *Glud1, Slc1a5*, *Slc7a5* and *Cpt1*, a rate-limiting step in fatty acid oxidation (**Fig. 5D and F-I)**. These data suggest that mTOR signaling governs a distinct transcriptional program during glucose recovery to enable the reactivation of previously stored metabolic and proliferative potential.

Collectively, our findings support a model in which environmental nutrient availability functions as an independent regulator of CD8+ T cell proliferative potential. Rather than division capacity being fixed by initial activating signals and decaying in a c-Myc-dependent manner, we demonstrate that nutrient-restricted T cells can preserve this potential over time. Upon nutrient recovery, CD8+ T cells are capable of rapidly increasing c-Myc expression, metabolic capacity, and reengaging mitogenic signaling to accelerate division kinetics. Notably, this occurs even in the absence of additional activating stimuli. This nutrient-responsive proliferative burst is not solely governed by c-Myc, as we find the mTOR pathway emerges as a dominant regulator through its unique control of metabolic gene expression in these cells. Thus, mitogenic and nutrient signaling independently orchestrate CD8+ T cell expansion capacity by converging on a shared anabolic program, yet each relies on distinct signaling arms of that pro-growth network to mediate its effects.

## Discussion

Cellular proliferation is an essential process required for organismal homeostasis, allowing tissues to renew, repair, and adapt to physiological demands. Uncontrolled proliferation can lead to oncogenic transformation, and cells tightly coordinate when and to what extent they divide in response to external cues^81,82^. In most mammalian systems, growth factor-driven signaling is the primary gatekeeper of cell cycle entry and progression, dictating a cell’s division capacity. This paradigm also applies to CD8+ T cells. Signals from the TCR, costimulation, and cytokines set the limit of clonal expansion through their control of key pro-growth signaling effectors, including MAPK-ERK, PI3K-AKT-mTOR, and c-Myc^38–41,51^. Here, we instead demonstrate that nutrient availability functions as an independent regulatory switch governing these pro-growth pathways. Beyond serving as a rate-limiting substrate, environmental glucose dynamically throttles CD8+ T cell expansion capacity. We find that CD8+ T cells activated under glucose restriction retain their mitogen-licensed division potential. Upon glucose restoration, T cells are able to fully realize this latent capability. This occurs in the absence of additional mitogenic input and results in the rapid elevation in metabolic capacity and pro-growth signaling. Notably, while glucose recovery leads to elevated c-Myc protein expression and ERK signaling, these factors are insufficient in the absence of mTOR activity, underscoring the unique role of mTOR signaling in this phenomenon. These findings support a model in which both mitogenic and metabolic inputs govern division potential in parallel through their distinct interactions with key anabolic signaling nodes. This enables cells to tune the rate at which they convert proliferative potential into cell division and align their mitogenically licensed program with nutrient availability. Altogether, our study offers a new perspective on the ability of CD8+ T cells to flexibly execute their host-protective function across an unpredictable landscape of metabolic environments.

In CD8+ T cells, the strength of the activating signal determines both the kinetics of quiescence exit and the magnitude of clonal expansion^33,38,74^. Using OT-I T cells stimulated with varying peptide affinity, antigen dose, or CD3/CD28 concentration, we show that mitogenic input controls the kinetics of activation, cell cycle entry, and survival, collectively shaping the expansion outcome, which is consistent with prior reports^38,51,59^. In contrast, we find that while glucose similarly affects CD8+ T cell expansion potential, it does so predominantly through its effects on cell cycle kinetics, with minimal impact on activation, survival, or even c-Myc expression. In this manner, both mitogenic signaling and nutrient availability cooperate to determine proliferative output, yet through distinct and underexplored mechanisms. Our findings address a key gap in the understanding of how these two critical inputs act in parallel circuits, instead of a linear hierarchy, to shape the expansion outcome. We show that restoring glucose to previously restricted cells, even several days after activation, not only reinstates but accelerates proliferation. This reveals that metabolic availability can dynamically preserve and re-engage proliferative potential independent of renewed mitogenic stimulation. Thus, metabolic cues can override temporal limits imposed by mitogenic signaling. Our work supports a model in which proliferative potential is cooperatively governed through nutrient- and mitogen-responsive axes, rather than a fixed property preprogrammed solely by receptor and growth factor signaling with nutrient uptake as a downstream consequence.

mTORC1 and ERK signaling are well-established drivers of early T cell activation, enabling both quiescence exit and cell cycle progression^33,42–44,49,83^. c-Myc functions downstream of these pathways as a key transcriptional regulator, and its expression has been proposed to act as a “division timer”, scaling with activating signal strength and gradually decaying to deplete proliferative potential^31,51,59,64^. Consistent with this model, we observe that canonical mTOR targets, including pS6K, pS6 and c-Myc, reach their peak levels within the first 24-48 hours of activation before diminishing over time. However, glucose restriction reveals an unanticipated divergence in this signaling network. Cells cultured under restricted conditions exhibit elevated pAKT (Ser473) and pERK, despite reduced mTORC1 activity and c-Myc expression. Upon glucose restoration, mTORC1 and c-Myc are rapidly reactivated, accompanied by further enhancement of ERK and mTORC2-AKT signaling. This coordinated resurgence is paralleled by pronounced metabolic remodeling, characterized by elevated mitochondrial respiration and glycolytic activity^63^. The resulting increase in ATP production likely supports the heightened bioenergetic demands of glucose-restored cells undergoing rapid proliferation. Together, these findings suggest that nutrient-limited CD8+ T cells preserve distinct signaling modules (mTORC2-AKT and ERK) that maintain proliferative readiness. Nutrient reexposure then re-engages a mTOR-driven growth program, highlighting how metabolic and mitogenic inputs can differentially modulate AKT, mTORC, c-Myc, and ERK activity to control CD8+ T cell expansion.

Once CD8+ T cells have fully entered the cell cycle, they are generally thought to become less reliant on mTOR activity for continued proliferation^42–44^. However, the heightened sensitivity of glucose-recovered cells to mTOR inhibition illustrates that mTOR’s role in cell cycle regulation is highly context dependent. These findings suggest a phase-specific function for mTOR after quiescence exit. Whereas mTOR is largely dispensable following early activation and during glucose restriction, it becomes essential to support the accelerated proliferation in glucose-recovered cells. This likely reflects that T cells distinctly require rapid metabolic reprogramming and biomass accumulation enforced by mTOR at inflection points in cell cycle kinetics, like quiescence exit and the glucose-throttled dynamics we studied. Indeed, it is well appreciated that T cells display distinct metabolic requirements for the cell cycle at different stages of activation and proliferation^49,56,84–86^. Further highlighting how distinct anabolic growth factors govern the unique demands of CD8+ T cell proliferation across contexts, we find mTOR inhibition does not blunt the glucose recovery-driven elevation in c-Myc levels. Conversely, though c-Myc is also nutrient sensitive, it plays a larger role in monotonic CD8+ T cell expansion through its temporal decay. We propose that the mTOR pathway acts as the essential throttling factor controlling cell cycle acceleration during metabolic transitions, whereas c-Myc governs the temporal progression toward cell cycle arrest, with both factors independently regulated by nutrient and mitogenic inputs. Together, this network both sets the initial expansion capacity and converts latent proliferative capacity into accelerated expansion when restricted cells encounter nutrient restoration.

These findings help revise the prevailing model in which proliferative potential is fixed at the time of activation and monotonically executed thereafter. We demonstrate that the metabolic environment dynamically paces the timing with which mitogenically licensed expansion capacity is stored or actualized, operating as a mitogen-independent circuit. Consequently, nutrient availability can temporally separate the establishment of proliferative potential from its deployment. In the context of adaptive immunity, CD8+ T cells initially receive their activating signals at sites like draining lymph nodes and subsequently seek out their target antigen in distal tissues. This split regulation of proliferative potential enables clonal T cell lineages to traverse dynamic nutrient environments while retaining the ability to undergo rapid expansion conferred in the lymph node. Accordingly, high-affinity clones are not irrevocably lost or hamstrung upon entering dysregulated tissue environments. Rather, they can later recover their affinity-driven expansion potential, allowing clonal selection to still identify the most fit clones regardless of the environmental path taken.

More broadly, nutrient-dependent preservation of proliferative capacity may operate in other dynamically cycling systems, including stem and progenitor cells, epithelial regeneration and cancer cells^87^. By uncoupling mitogenic commitment from metabolic execution, our work reframes nutrient availability as an active regulator, not just a permissive substrate, of expansion fate. This highlights metabolic plasticity as a key determinant of the long-term proliferation outcomes of adaptive immunity^88^. Moving forward, it will be key to understand how T cells sense and store proliferative potential during nutrient stress and what factors trigger its release. Uncovering these mechanisms will offer strategies to therapeutically harness the stored CD8+ T cell expansion capacity during infection and cancer or restrict it in autoimmune disease and transplantation.

## Materials and Methods

### Mice

OT-I (Jax #003831) mice were purchased from Jackson Laboratories and maintained in house. All mice were bred and housed under specific pathogen-free conditions with a 12-hour on/off light cycle at the Children’s Hospital of Philadelphia. All experiments were performed with female or male mice aged between 6 to 10 weeks of age. All experiments were performed in accordance with the Institutional Animal Care and Use Committee of the Children’s Hospital of Philadelphia (IAC 21–001325).

### CD8+ T cell Isolation and Culture

Murine CD8+ T cells were isolated from spleen and lymph nodes, following ACK-lysis, by negative selection using an EasySep™ Mouse CD8+ T Cell Isolation Kit (STEMCELL Technologies, #19853). For experiment to test TCR strength and affinity, OT-I T cells were stimulated for 24 hours with either plate-bound anti-CD3ε/CD28 activation at 5µg/mL or 1µg/mL: anti-CD3ε (clone 145-2C11, BioLegend, #100360, RRID:AB_2800555) and anti-CD28 antibodies (clone 37.51, BioLegend, #102122, RRID:AB_11147170) or with either SIINFEKL (N4) (Anaspec, #AS-60193-1) or SIIGFEKL (G4) (Anaspec, #AS-64384) at 1 μg/mL or 0.01 μg/mL and 2.5µg/mL anti-CD28 and rIL-2 (5ng/mL, BioLegend, #575406). After 24 hours of activation, cells were removed from stimulation and seeded at 5 x 10^4^ cells/well, then cultured in 96-well plates.

T cell stimulation and culture was performed in supplemented lymphocyte culture medium: Roswell Park Memorial Institute (RPMI) 1640 Medium (Gibco, #11875093) or No Glucose = RPMI 1640 Medium, no glucose (Gibco, #11879020) with 10% dialyzed fetal bovine serum (Corning, #MT35071CV), additional 2mM L-glutamine (Gibco, #25030081), 1mM sodium pyruvate (Gibco, #11360070), 25mM HEPES (Gibco, #15630080), 1x penicillin/streptomycin (Gibco, #15140122), 55μM 2-mercaptoethanol (Gibco, #21985023). For experiments involving activated and maintained T cells in different glucose conditions, the following concentrations were used: 0.1 mM, 2.5 mM, 5 mM and 10 mM glucose (Fisher, #BP350500). All cells were cultured in a CO_2_ incubator at 37°C with 5% CO_2_. For all experiments using pharmacological reagents, the following doses were used unless otherwise noted: Torin1 (50nM, Cayman Chemical, #10997) and Rapamycin (50nM, Sigma, #R0395).

### Flow cytometry

All flow cytometry was performed on a CytoFLEX LX or CytoFLEX S cytometer (Beckman Coulter) or 5-laser Cytek Aurora (Cytek Biosciences). Data were analyzed using FlowJo software (Becton Dickinson). Cell division was measured by labelling cells with CellTrace Violet (Invitrogen, #C34557) before activation and evaluating proliferation at the indicated time points. Staining was performed with combinations of the following antibodies: anti-CD8 (1:300; clone 53-6.7, Thermo Fisher Scientific), anti-CD69 (1:300, clone H1.2F3, Thermo Fisher Scientific), Ki-67 (1:600; clone SolA15, Thermo Fisher Scientific), anti-c-Myc (1:400; D84C12, Cell Signaling Technology) with secondary anti-rabbit IgG conjugated to Alexa Fluor 488 (1:750, A11008, Thermo Fisher Scientific). Viability was assessed using eBioscience Fixable Viability Dye eFluor 780 (1:1500, Thermo Fisher Scientific, #65-0865-18). Cells were first stained with surface antibodies and viability dye diluted in PBS for 30 min on ice, then fixed with 1x Fixation/Permeabilization Buffer for 30 min on ice (eBioscience™ Foxp3 / Transcription Factor Staining Buffer Set, Invitrogen, #00-5523-00), and lastly stained with intracellular antibodies diluted in 1x Permeabilization Buffer (eBioscience™ Foxp3 / Transcription Factor Staining Buffer Set, Invitrogen, #00-5523-00) for 30 min-1h on ice. Samples were run in technical replicates.

For the EdU incorporation assay, 10 x 10^4^ CD8+ T cells were cultured as indicated and labeled with 5 mM EdU for 4 hours of culture. EdU incorporation was detected using the Click-iT™ Plus EdU Alexa Fluor™ 488 Flow Cytometry Assay Kit (Invitrogen, C10632), according to manufacturer’s instructions. DNA content was measured by DAPI to determine cell cycle state.

For cell counting, a known number of precision count beads (BioLegend, #424902) was added to samples immediately before analysis, and the ratio of beads to live cells was used to calculate the absolute cell number in each sample.

### Mitochondrial staining

For mitochondrial staining, 10 x 10^4^ CD8+ T cells were stained with 50nM MitoTracker Green (MTG) (Thermo Fisher Scientific, #M7514), 100nM Tetramethylrhodamine, Ethyl Ester, Perchlorate (TMRE) (Thermo Fisher Scientific, #T668), anti-CD8 (1:300; clone 53-6.7, Thermo Fisher Scientific) and eBioscience Fixable Viability Dye eFluor 780 (1:1500, Thermo Fisher Scientific, #65-0865-18) for 30 min at 37°C. After this, cells were washed twice and resuspended in PBS. Cells were immediately acquired, and mitochondrial readouts were gated on live CD8+ T cells.

### RT-qPCR

RNA was extracted using the Zymo Quick-RNA MicroPrep Kit (Zymo Research, #R1050) with DNase I treatment, as per the manufacturer’s protocol. cDNA was synthesized with 1µg of purified RNA using Thermo Maxima H Minus Reverse Transcriptase (Thermo Scientific, #EP0751) and following the manufacturer’s instructions using 25 pmol oligo(dT)_18_ and 25 pmol random hexamer primers. The reverse transcribed reaction was incubated for 10 min at 25°C, 15 min at 50°C, and then 5 min at 85°C. cDNA was diluted 1:10 ratio with molecular grade water and quantified via real-time quantitative polymerase chain reaction (RT-qPCR) with PowerTrack SYBR Green Master Mix (Thermo Scientific, #A46110) on a Bio-Rad CFX384 real-time qPCR system. Quantified transcripts were normalized to *Rps18*, and data from all samples were plotted as relative expression versus the control group in DMSO.

*Hif1a* F – ACTTCTGGATGCCGGTGGTCT

*Hif1a* R – TGTCGCCGTCATCTGTTAGCAC

*Slc2a1* F – CACCACACTCACCACGCTTTG

*Slc2a1* R – ATGGAGTTCCGCCTGCCAA

*Slc1a5* F – TTTGGTGTGGCTCTGCGGAA

*Slc1a5* R – CCAACGGGTGCGTACCACATA

*Slc7a5* F – AGCTGTGGCTGTGGACTTCG

*Slc7a5* R – CCCATTGACAGAGCCGAAGCA

*Hk2* F – CAACTCCGGATGGGACAGAACA

*Hk2* R - ACCCTTACTCGGAGCACACG

*AldoA* F – TGTTCTGCCTTACAGATCCTGGAC

*AdolA* R – TGGCAGTGCTTTCCGGTCTT

*Eno1* F - CTTCATGGGGAAGGGTGTCTCA

*Eno1* R – TGTCAATCTTCTCTTGCTCCACAAC

*Pgk1* F – ATCTGCCACAGAAGGCTGGTG

*Pgk1* R – TGCAACTTTAGCGCCTCCCAA

*Ldha* F - TCGTGCACTAGCGGTCTCAA

*Ldha* R – GTTCTGGGGAGCCTGCTCTT

*Glud1* F – AAGTGCGCTGTGGTCGATGTA

*Glud1* R – CAGCTCCATAGTGAACCTCCGT

*Cpt1a* F – TGTACGCTCCTGCACTACGG

*Cpt1a* R - TGAACCTCTGCTCTGCCGTT

### Western blot

Cells were pelleted, washed in PBS and lysed in ice-cold RIPA buffer (pH = 8.0) containing 50 mM Tris, 1 mM EDTA, 150 mM NaCl, 1% NP-40, 0.5% Na-deoxycholate, 0.1% SDS and supplemented with 10 mM NaF, 1 mM Na_3_VO_4_, and EDTA-free Protease Inhibitor Cocktail (Roche, #11836170001). Protein lysate concentration was quantified via Bradford assay (Bio-Rad, #5000006), using bovine serum albumin as a standard. 20-25µg protein lysate was mixed with reducing sample buffer (Boston BioProducts, #BP-111R), and samples were separated by SDS-PAGE on Any kD TGX Precast Protein Gels (BioRad, #456-8125), followed by transfer using the Trans-Blot Turbo Transfer kit (BioRad, #1704274) on the Turbo setting. Membranes were blocked with 5% dry nonfat milk in tris-buffered saline with 0.1% Tween 20 (TBS-T) for 60 min. Primary antibodies were diluted in TBS with 1% Casein (BioRad, #1610782) and incubated with membranes overnight at 4°C with rocking. Three TBS-T washes of 5 minutes each were performed before and after 1-hour incubation with secondary antibodies (diluted in 5% dry nonfat milk in TBS-T) at room temperature with rocking. Membranes were visualized with SuperSignal West Pico PLUS (Thermo Scientific, #34580) or Femto Chemiluminescent Substrate (Thermo Scientific, #34095) using the ChemiDoc MP Imaging System (BioRad). Blots were stripped using Restore Western Blot Stripping Buffer (Thermo Scientific, #21059) after visualization of phospho-epitopes, before incubating with total antibody. Antibodies used: Phospho-S6K (1:1000, 108D2 Thr^389^, Cell Signaling Technology), S6K (1:1000, 49D7, Cell Signaling Technology), Phospho-S6 (1:2000, D57.2.2E Ser^235/236^, Cell Signaling Technology), S6 (1:5000, 5G10, Cell Signaling Technology), c-Myc (1:1000, D84C12, Cell Signaling Technology), Phospho-AKT (1:1000, D9E Ser^473^, Cell Signaling Technology), AKT (1:1000, C67E7, Cell Signaling Technology), Phospho-ERK1/2 (1:1000, D13.14.4E Thr^202^/Tyr^204^, Cell Signaling Technology), EKR/12 (1:1000, #9102, Cell Signaling Technology), Phospho-4E-BP1 (1:1000, 236B4 Thr^37/46^, Cell Signaling Technology), 4E-BP1 (1:1000, 53H11, Cell Signaling Technology), Actin-hFAB rhodamine (1:2000, BioRad).

### Seahorse analysis

For Seahorse assay, 1.25 x 10^5^ cells were plated per well in a 96-well Seahorse assay plate precoated with Cell-Tak (Corning, #354240). For the mitochondrial stress test assay, cells were cultured in Seahorse XF RPMI medium (Agilent, #103576-100) supplemented with 10mM glucose, 2mM L-glutamine, and 1mM sodium pyruvate and treated with oligomycin (1.5 μM, Sigma-Aldrich, #O4876), BAM15 (2.5 μM, Cayman Chemical, #17811), and rotenone (0.5 μM, Cayman Chemical, #13995-1)/antimycin A (0.5 μM, Sigma-Aldrich, #A8674) respectively at indicated time points. For the glycolysis stress test assay, cells were cultured in glucose-free Seahorse XF RPMI (Agilent, #103576-100) supplemented with 2mM L-glutamine and 1mM sodium pyruvate prior to assay and treated with 10 mM glucose (Fisher, #BP350500), oligomycin (1 μM, Sigma-Aldrich, #O4876), and 2-deoxyglucose (50mM, Sigma-Aldrich, #D0051) respectively at indicated time points. Data analysis and transformation were performed using Wave software. ATP production from glycolysis was inferred from the Proton Efflux Rate (PER) measurements. Mitochondrial ATP production was calculated by subtracting the oxygen consumption rate after oligomycin injection from the baseline measurement to obtain OCR_ATP_, which was then converted to ATP production using the established stoichiometric relationship^61,62,89^.

### Statistical analysis

Statistical significance was calculated using the tests indicated in each figure legend, performed in GraphPad Prism software. All the error bars represent the standard error of the mean (SEM). We considered results with a p-value less than 0.05 (p< 0.05) to be statistically significant. * = p < 0.05, ** = p < 0.01, *** = p < 0.001, and **** = p < 0.0001. All experiments were repeated at least three times across independent experiments.

## Acknowledgments

We thank the Children’s Hospital Flow Cytometry Core for providing support and instrumentation, and all members of the Bailis Laboratory for feedback and support. The figure schematic was created in https://BioRender.com.

---

**Supplement Figure 1.**
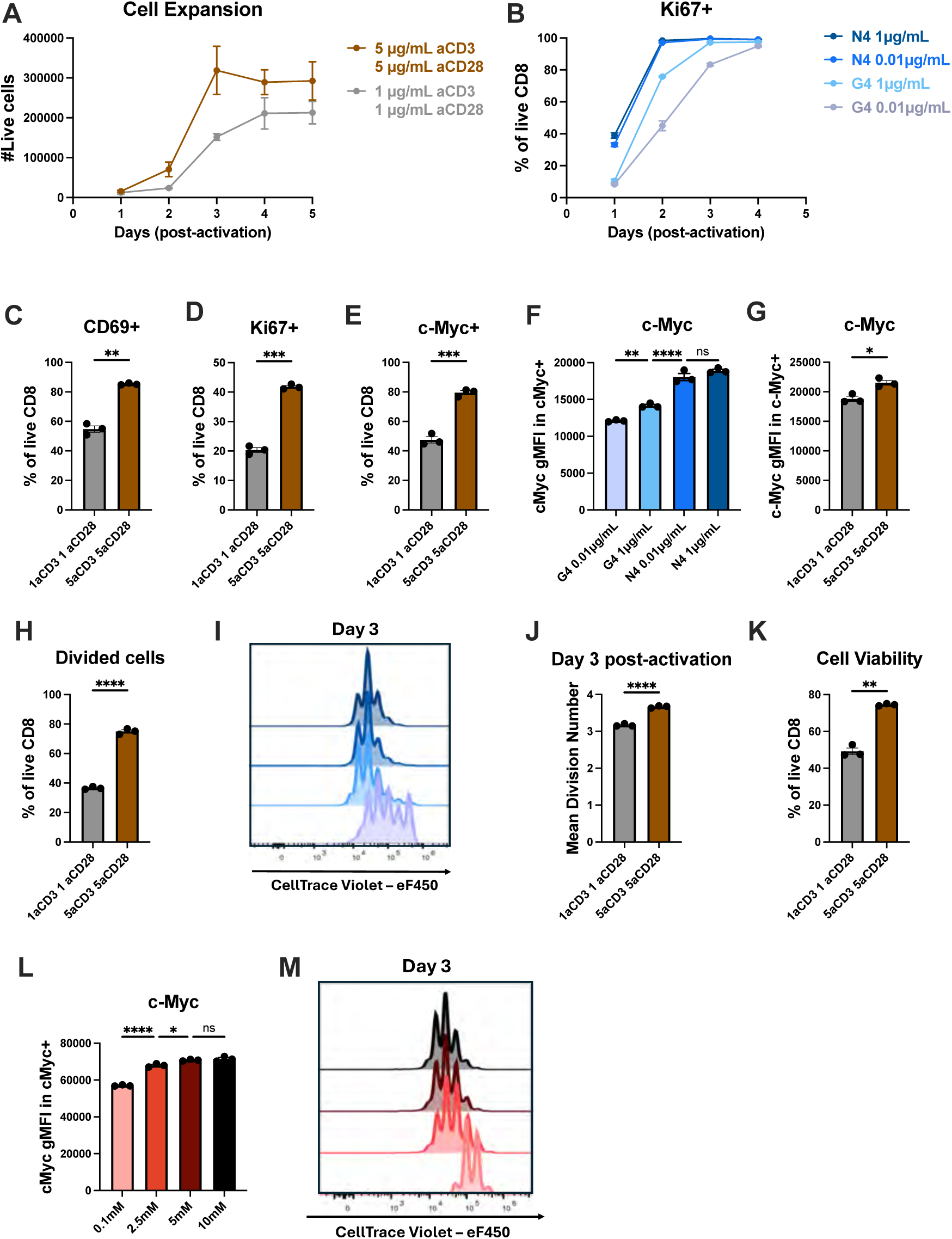
CD8+ T cell proliferation is controlled by both the strength of TCR stimulation and environmental nutrients. CD8+ T cells were purified from OT-I spleen and lymph nodes that had been stimulated for 24 hours with either 5µg/mL or 1µg/mL of anti-CD3/CD28. (**A**) Quantification of number of live CD8+ T cells activated by different anti-CD3/CD28 concentrations. (**B**) Frequency of CD8+ T cells entering cell cycle (Ki67+) over 4 days after activation. (**C**) Frequency of CD8+ T cells expressing activation marker CD69 24 hours post-activation. (**D**) Frequency of CD8+ T cells entering cell cycle (Ki67+) 24 hours post-activation. (**E**) Frequency of CD8+ T cells expressing c-Myc 24 hours post-activation. (**F**) c-Myc geometric mean fluorescence intensity (gMFI) in CD8+ T cells expressing c-Myc 24 hours post-activation. (**G**) c-Myc gMFI in CD8+ T cells expressing c-Myc 24 hours post-activation. (**H**) Frequency of divided cells (≥ 1 division) on day 2 post-activation. (**I**) Representative histogram of CellTrace Violet (CTV) on day 3 post-activation. (**J**) Mean division number on day 3 post-activation. (**K**) Frequency of live CD8+ T cells 24 hours post-activation. (**L**) c-Myc gMFI in CD8+ T cells expressing c-Myc 24 hours post-activation. (**M)** Representative histogram of CTV on day 3 post-activation. All error bars are representative of 3 technical replicates. Statistical significance in (C, D, E, G, H, J, and K) was calculated using Welch’s t-test, and statistical significance in (F and L) was calculated using one-way ANOVA with multiple comparisons and Tukey’s correction.

**Supplement Figure 2.**
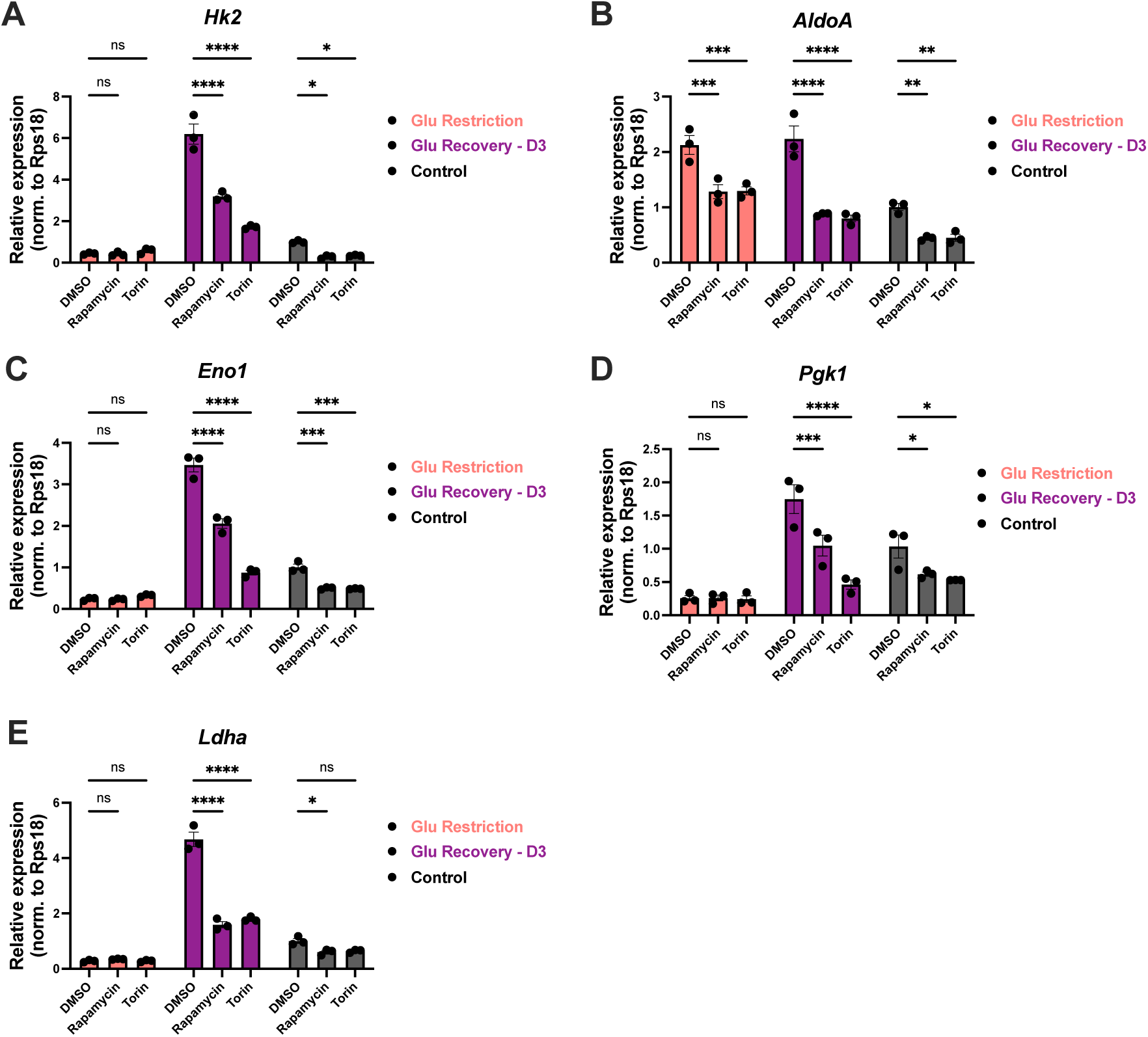
mTORC signaling is essential for increased transcript abundance of key glycolytic enzymes. On day 3 post-activation, cells in their respective media conditions were treated with DMSO, rapamycin (50nM), or Torin 1 (50nM), and samples were collected on day 4 (24-hour treatment). Relative mRNA expression of key glycolytic enzymes (**A**) hexokinase 2 (*Hk2*), (**B**) aldolase (*AldolA*), (**C**) enolase 1 (*Eno1*), (**D**) phosphoglycerate kinase 1 (*Pgk1*), (**E**) lactate dehydrogenase A (*Ldha*). All error bars are representative of 3 technical replicates. Statistical significance was calculated using two-way ANOVA with multiple comparisons and Tukey’s correction.

